# Neuroanatomical correlates of social intelligence measured by the Guilford test

**DOI:** 10.1101/2020.12.03.409466

**Authors:** A Myznikov, M Zheltyakova, A Korotkov, M Kireev, R Masharipov, O.Dz. Jagmurov, U Habel, M Votinov

## Abstract

Social interactions are a crucial aspect of human behaviour. Numerous neurophysiological studies have focused on socio-cognitive processes associated with the so-called theory of mind – the ability to attribute mental states to oneself and others. Theory of mind is closely related to social intelligence defined as a set of abilities that facilitate effective social interactions. Social intelligence encompasses multiple theory of mind components and can be measured by the Four Factor Test of Social Intelligence (the Guilford-Sullivan test). However, it is unclear whether the differences in social intelligence are reflected in structural brain differences. During the experiment, 48 healthy right-handed individuals completed the Guilford-Sullivan test. T1-weighted structural MRI images were obtained for all participants. Voxel-based morphometry analysis was performed to reveal grey matter volume differences between the two groups (24 subjects in each) – with high social intelligence scores and with low social intelligence scores, respectively. Participants with high social intelligence scores had larger grey matter volumes of the bilateral caudate, left insula, left inferior parietal lobule, inferior temporal gyrus, and middle occipital gyrus. Only the cluster in the caudate nuclei survived a cluster-level FWE correction for multiple comparisons. The obtained results suggest caudate nucleus involvement in the neural system of socio-cognitive processes, reflected by its structural characteristics.

## Introduction

Social interactions are a crucial part of everyday life. There is increasing support for so-called “social intelligence,” which is dissociable from general intelligence. Social intelligence is defined as a set of human abilities that facilitates effective interaction with other people, including the ability to infer the mental and emotional states of others, to understand their intentions, and to predict their behaviour (Moss and Hunt, 1927; Thorndike, 1920; Vernon, 1933). Evidence points to an association of this skill with socio-cognitive processes of the so-called theory of mind (TOM), the ability of a person to attribute mental states (for example, thoughts, beliefs, and intentions) to oneself and to others (Premack and Woodruff, 1978).

TOM processes are the focus of many neurophysiological studies, and various experimental tasks have been used to define their neural correlates. A classic example of such a task is making predictions or assumptions based on a story in which a protagonist has a false belief (Dodell-Feder et al., 2011; Saxe et al., 2006). Other studies used different stimulus modalities, in particular, cartoons (Völlm et al., 2006), photographs (Canessa et al., 2012; de Lange et al., 2008), videos (Boccadoro et al., 2019; Wolf et al., 2010), and animations (Gobbini et al., 2007), representing protagonists’ actions or social interactions with the task, either to passively view the stimuli or to make assumptions based on the information presented. Another test measuring TOM ability is Reading the Mind in the Eyes Test (RMET), where a participant is presented with photographs of the eye region and is asked to decide which word better describes what a person in the picture is thinking or feeling (Baron ‐ Cohen et al., 2001). Some studies also use interactive game designs, such as the prisoner’s dilemma game for engaging TOM (Kircher et al., 2009). According to a meta-analytic study, neurophysiological correlates of TOM processes reported in different papers can vary as a result of task-related activation because studies focus on different aspects of TOM (e.g., implicit versus explicit, cognitive versus affective, visual versus verbal TOM) (Molenberghs et al., 2016).

The issues mentioned above can be potentially improved by using more sophisticated psychological tasks that engage multiple components of TOM. An appropriate example is the standardised J. Guilford and M. O’Sullivan Four Factor Test of Social Intelligence (Guilford-Sullivan test) that aims to give a composite evaluation of a person’s social intelligence isolated from general intelligence. This test is based on Guilford’s structure-of-intellect model. According to this model, social intelligence includes 30 different abilities, four of which are quantitatively measured in four subtests of the Guilford-Sullivan test (O’Sullivan, 1965). The subtests estimate (1) the ability to group other people’s mental states based on similarity – Expression Grouping subtest, (2) the ability to interpret sequences of social behaviour – Missing Cartoons subtest, (3) the ability to respond flexibly in interpreting changes in social behaviour – Cartoon Prediction subtest, and (4) the ability to predict what will happen in an interpersonal situation – Social Translations subtest. These subtests include different stimuli modalities (verbal and visual) and various tasks (grouping, predicting, and interpreting), measuring several components of socio-cognitive processes. They are closely related to tasks frequently utilized to measure TOM. For example, the Expression Grouping subtest is closely associated with RMET, while Missing Cartoons and Cartoon Prediction subtests are widely and independently used to estimate TOM (Brunet et al., 2000; Gallagher et al., 2000; Völlm et al., 2006). To summarize, the Guilford-Sullivan test encompasses four measurements obtained by different TOM-related tasks. Thus, it can be a good psychological scale for studying the neural correlates of socio-cognitive processes.

Several clinical neuroimaging studies used the Guilford-Sullivan test to assess its correlation with neuroanatomical characteristics based on VBM data. It was demonstrated that impairments in this test's performance in first-episode psychosis were significantly correlated with reduced grey matter density in the left middle frontal gyrus, the right supplementary motor cortex, the left superior temporal gyrus, and the left inferior parietal lobule (Bertrand et al., 2008). Along with that, Cartoon Prediction subtest performance in patients with social and executive disorders, in the case of frontotemporal dementia, was correlated with GMV in the orbital frontal, superior temporal, visual association, and posterior cingulate regions of the right hemisphere (Eslinger et al., 2007). Although the Guilford-Sullivan test was only used to study pathologies, mentioned brain structures can also be associated with social intelligence in healthy people.

At the same time social intelligence is a measure of socio-cognitive abilities. And a broader list of brain regions is reported in association with social cognition processes. This evidence is gained partly from analyses of the human brain’s structural characteristics using voxel-based morphometry analysis (VBM analysis). In particular, performance in RMET was positively correlated with the grey matter volume (GMV) of the dorsomedial PFC, inferior parietal lobule (TPJ), and precuneus in the left hemisphere (Sato et al., 2016) as well as with the volume of the caudate nucleus and putamen (Warrier et al., 2018). In another study, performance in a similar facial expression recognition test was positively correlated with the volume of the right inferior frontal gyrus (IFG) (Cabinio et al., 2015). In a task providing conditions for spontaneous social interactions, a significant positive correlation was shown between the participant’s performance and cortical thickness in the medial PFC, right IFG, and right TPJ (Rice and Redcay, 2013).

Along with structural findings, functional studies expanded the set of brain regions attributed to socio-cognitive processes. For example, the striatum was implicated in different aspects of social behaviour, such as social reward and learning about others’ preferences, according to the results of human and animal studies (Báez-Mendoza and Schultz, 2013). In addition, the insula, as a part of the limbic cortex, was reported to be involved in socio-emotional processes, decision making in a social context, and social pain processing (Eisenberger, 2012; Lamm and Singer, 2010; Menon and Uddin, 2010; Uddin et al., 2017). In a meta-analysis of 350 fMRI studies on social cognition, Van Overwalle et al. found activation of the cerebellum during abstract mentalizing (Van Overwalle et al., 2014). Named structural and functional studies did not target social intelligence directly, but their results can potentially include its neural correlates, because it falls within the definition of socio-cognitive processes.

It was mentioned that the Guilford-Sullivan test includes tasks used to measure TOM abilities. Therefore, social intelligence can be associated with the neural correlates similar to those of TOM. There is evidence of involvement of several brain regions in TOM-specific socio-cognitive processes. In the meta-analysis, Molenberg and colleagues (2016) defined core areas of the TOM neural system repeatedly engaged in all types of tasks found in the TOM-related research literature: the medial prefrontal cortex (PFC) and bilateral temporoparietal junction (TPJ). Along with the precuneus and the right superior temporal sulcus, these areas were also reported in a study that utilized a task with stories about false beliefs on a substantial sample (N = 462) of individuals (Dufour et al., 2013). Named areas are believed to constitute the TOM neural system and, as a result can be related to social intelligence.

Thus, studies applying the Guilford-Sullivan test for identifying neural correlates of socio-cognitive processes are scarce, and their results are controversial. To our knowledge, there is no study on the association between GMV and the level of social intelligence in a healthy population. Besides, the results of studies using classical TOM tasks, together with other structural and functional analyses, cumulatively demonstrate that, depending on the type of experimental task, different brain areas can be attributed to the TOM neural system. Thus, this study aims to confirm and to expand the list of neuroanatomical correlates of socio-cognitive processes using the Guilford-Sullivan test and VBM analysis. Based on prior studies, it can be hypothesized that this test’s performance for healthy people will correlate with the GMV in previously defined nodes of the TOM neural system in the medial PFC, the TPJ in both hemispheres, the precuneus and the right superior temporal sulcus (Dufour et al., 2013; Molenberghs et al., 2016). In addition, it can be assumed that the level of social intelligence will correlate with the GMV in regions reported in clinical studies that utilized Guilford-Sullivan test, such as cingulate cortex and orbitofrontal cortex.

## Methods

A total of 48 healthy right-handed volunteers (32 women and 16 men) participated in the study. All participants were 24,9±5,5 years old, with no history of neurological or psychological disorders and no contraindications for magnetic resonance imaging. All subjects provided written informed consent prior to the study. All procedures were conducted in accordance with the Declaration of Helsinki and were approved by the Ethics Committee of the N.P. Bechtereva Institute of the Human Brain, Russian Academy of Sciences.

### Social intelligence testing

The Russian adaptation of the four-factor test of social intelligence, developed by J. Guilford and M. Sullivan (Guilford-Sullivan test), was used to measure the level of social intelligence (Mихайлова E.C, 2001). This test consists of four subtests: 1) Cartoon Predictions, 2) Expression Grouping, 3) Social Translations, and 4) Missing Cartoons (see Figure 1). In the first subtest (Cartoon Predictions), participants need to choose one out of three cartoons, which appropriately continues the suggested situation. The task in the expression grouping subtest is to find the facial expression that best fits a group of three other expressions. In the social translations subtest, a statement between a pair of people in certain social situations is presented; participants need to choose one out of three situations in which a suggested statement has a different meaning. Within the Missing Cartoons subtest, participants’ task is to choose one out of four cartoons, which completes the suggested scenario in the right way. The first subtest consisted of 14 tasks, while the second, third, and fourth subtests consisted of 15, 12, and 14 tasks, respectively. Considering that these subtests vary in the type of activity and can be expressed to varying degrees in different subjects, a cumulative measure was utilised to obtain the balanced individual characteristic of social intelligence rather than to investigate its particular components.

**Figure 1.**
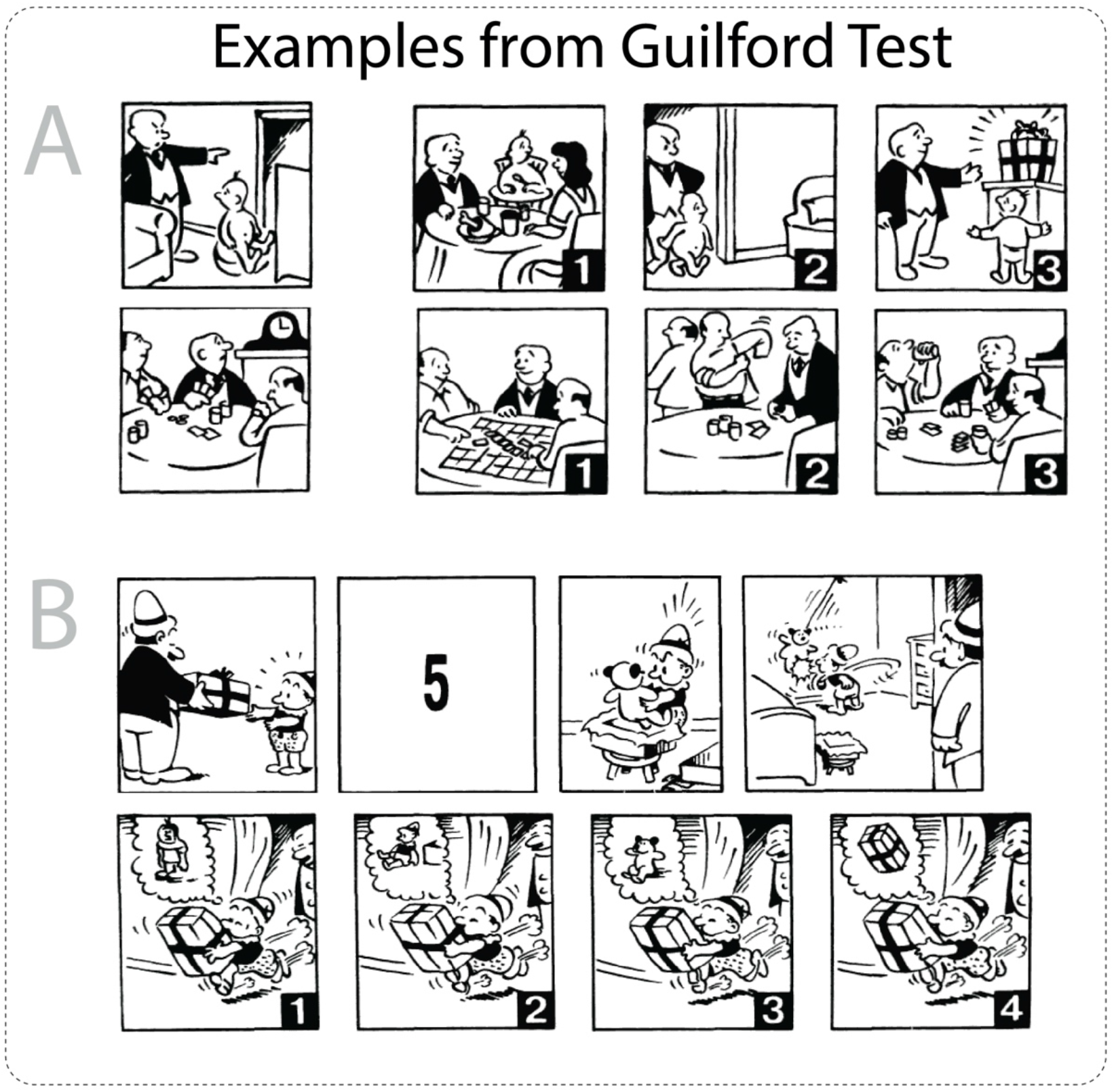
Examples of measurements in the Guilford-Sullivan test. Task A represents the Cartoon Predictions subtest, where participants must select one of three cartoons that most appropriately describes the outcome of the suggested situation. Task B represents the Missing Cartoons subtest, where participants are required to choose one of four cartoons, which correctly fills the suggested sequence of cartoons.

Raw scores for all subtests were summed and then transformed to a standard test score (see Table 1), which ranged from 1 (low level of social intelligence) to 5 (high level of social intelligence). Standard scores for every subject were used as a criterion of group formation for the VBM analysis. Volunteers were divided into two groups (24 subjects in each): values equal to or below the score of 3 were considered low social intelligence, while values above the score of 3 were considered high social intelligence (see Table 2). Groups did not differ significantly in age, gender, or education.

**Table 1.** Conversion of raw Guilford-Sullivan test scores into standard scores

| **Raw score** | **Standard score** |
| --- | --- |
| **0-12** | **1** |
| **13-26** | **2** |
| **27-37** | **3** |
| **38-46** | **4** |
| **47-55** | **5** |

**Table 2.**
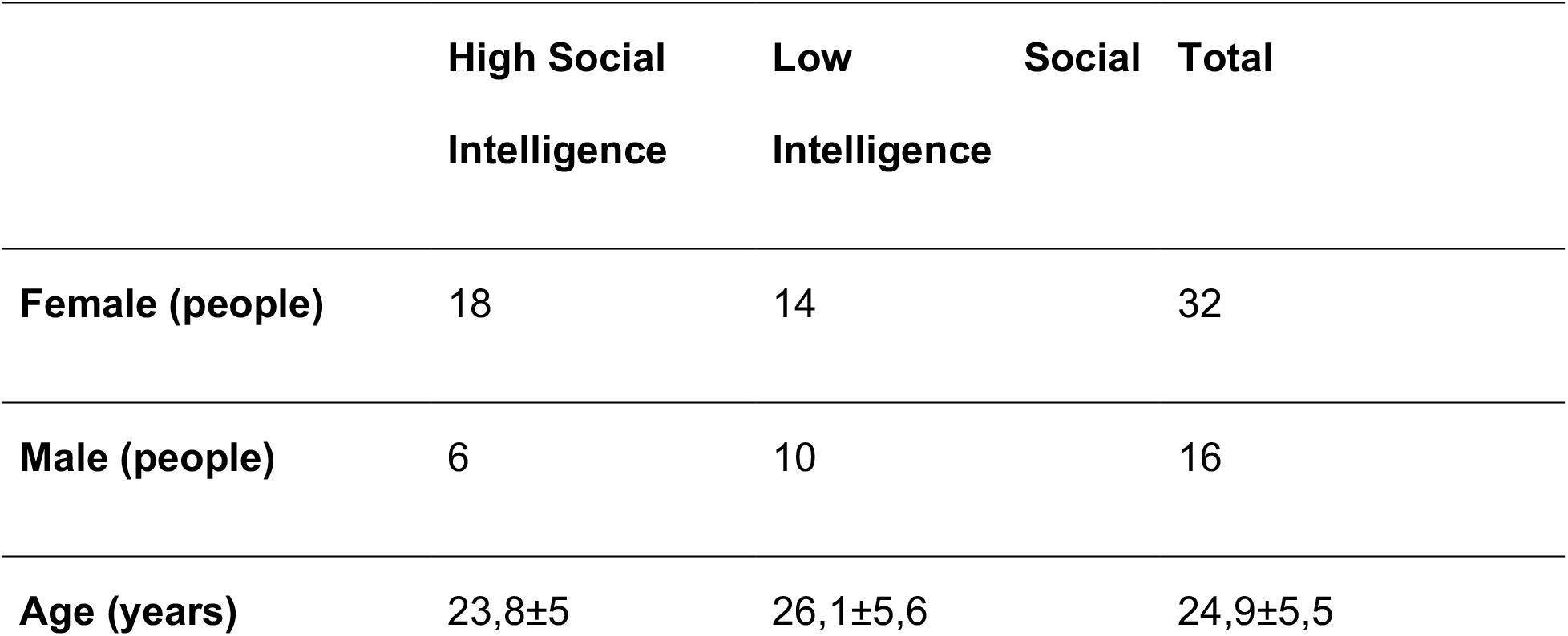
The demographics and the Guilford-Sullivan test scores

**Table 3.** Clusters of the grey matter volume differences associated with both [High Social Intelligence > Low Social Intelligence] and [Low Social Intelligence > High Social Intelligence] contrasts, minimal cluster size (k = 30)

| Region (L - left, R - right) | cluster size (k) | cluster-level p FWE-corrected | MNI coordinates |  |  |
| --- | --- | --- | --- | --- | --- |
|  |  |  | X | y | z |
| **High Social Intelligence > Low Social Intelligence** |  |  |  |  |  |
| R Caudate, L Caudate | 824 | 0.012 | 11 | 6 | 9 |
| L Posterior Insula | 183 | 0.518 | -42 | -14 | 8 |
| L Inferior Parietal Lobule | 113 | 0.753 | -41 | -54 | 57 |
| L Inferior Temporal Gyrus | 115 | 0.746 | -48 | -57 | -8 |
| L Middle Occipital Gyrus | 68 | 0.899 | -48 | -83 | 9 |
| L Precentral Gyrus | 66 | 0.904 | -57 | -6 | 15 |
| **Low Social Intelligence > High Social Intelligence** |  |  |  |  |  |
| L Middle frontal gyrus | 67 | 0.901 | -36 | 38 | 42 |
| L Inferior temporal gyrus | 185 | 0.512 | -38 | -5 | -41 |

### Data acquisition and quality control

Magnetic resonance imaging was performed using a 3 Tesla Philips Achieva (Philips Medical Systems, Best, The Netherlands). Structural images were acquired using a T1-weighted pulse sequence (T1W-3D-FFE; repetition time [TR] = 2.5 ms; TE = 3.1 ms; 30◦ flip angle), recording 130 axial slices (field of view [FOV] = 240 × 240 mm; 256 × 256 scan matrix) of 0.94 mm thickness. All MRI scans were inspected for image artefacts and incidental brain abnormalities. All subjects were included in the study.

### Voxel-based morphometry analysis (VBM-analysis)

The VBM analysis of structural data was performed with Statistical Parametric Mapping software (SPM12, Wellcome Department of Imaging Neuroscience, London, UK (www.fil.ion.ucl.ac.uk/spm) and the Computational Anatomy Toolbox 12 (CAT12; *http://dbm.neuro.uni-jena.de/cat.html*) running in MATLAB (MathWorks, Natick, MA). All structural data were manually reoriented to place their native-space origin at the anterior commissure. Then, the default parameters of the CAT12 toolbox were used. Images were corrected for magnetic field inhomogeneities and segmented into grey matter, white matter, and cerebrospinal fluid. Normalisation to MNI space using the DARTEL (Diffeomorphic Anatomical Registration Through Exponentiated Lie algebra) algorithm to a 1.5 mm isotropic adult template provided by the CAT12 toolbox was performed for segmented grey matter data. Finally, the grey matter segments were smoothed with a Gaussian smoothing kernel of 8 mm. The CAT12 toolbox provides an automated quality check protocol. Therefore, quality check control for all structural data was performed to obtain so-called image quality rating (IQR) scores, which were later used as an additional covariate in the statistical analysis. In addition, total intracranial volumes (TIVs) were calculated to be used as a covariate.

### Statistical analysis

The statistical analysis was performed for two groups of subjects: 1) those with a high social intelligence score (>3) and 2) those with a low social intelligence score (≤3). For the VBM analysis, we included the following confounders (as covariates), which can affect VBM results: sex (male/female), age, TIV, and IQR scores. The two-sample t-test was performed to test the hypothesized differences in the GMV. Statistical parametric maps were created with the uncorrected p-value (<0.001) and a subsequent cluster-level family-wise error (FWE) correction with p<0.05. The SPM results were visualized using the MRIcron toolbox (https://www.nitrc.org/projects/mricron).

## Results

The VBM analysis with the voxel-wise uncorrected p-value (<0.001) threshold for the “High Social Intelligence > Low Social Intelligence” contrast revealed significant GMV differences in the bilateral caudate nucleus, left insula, left inferior temporal gyrus, and left angular gyrus (see Table 2 and Figure 2). In the case of the “Low Social Intelligence > High Social Intelligence” contrast, two clusters in the left inferior temporal gyrus and left middle frontal gyrus were revealed (Table 2). After applying a stricter cluster-level FWE-corrected (p<0.05) threshold, only GMV differences in the bilateral caudate nucleus survived for the “High Social Intelligence > Low Social Intelligence” contrast (Figure 3), and nothing survived for the “Low Social Intelligence > High Social Intelligence” contrast.

**Figure 2.**
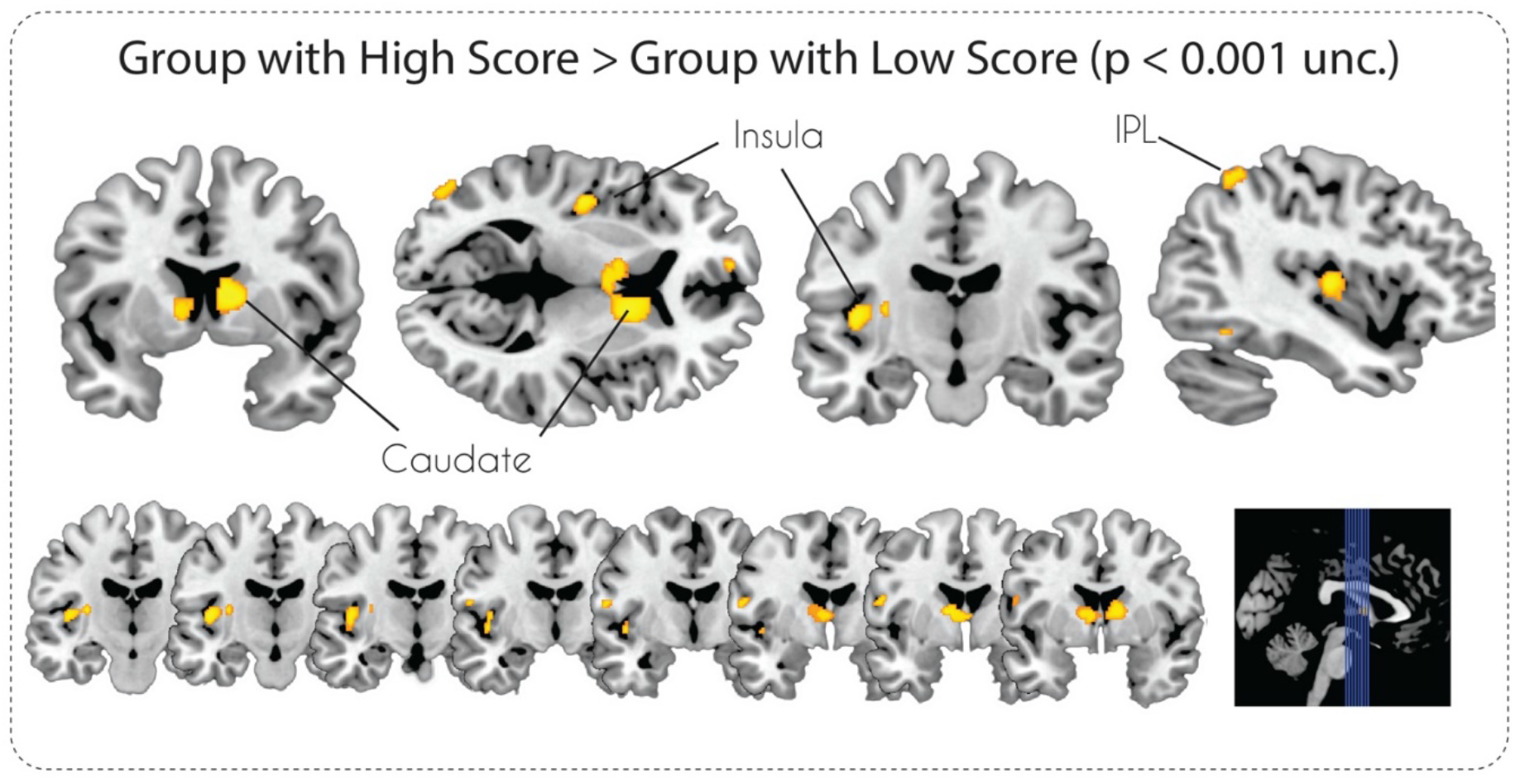
Statistical parametric maps of grey matter volume differences in subjects with High and Low Social Intelligence at p < 0.001, uncorrected.

**Figure 3.**
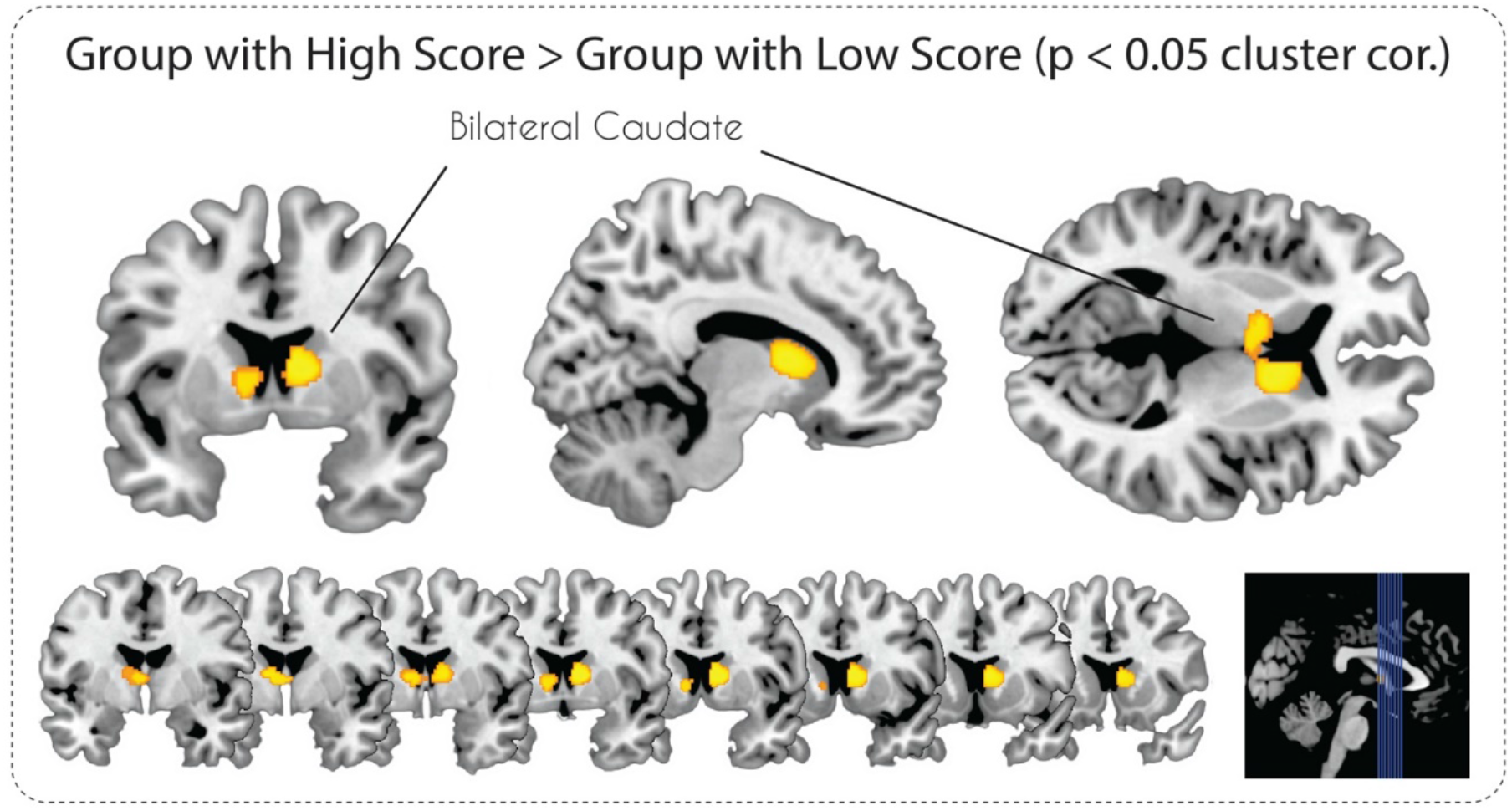
Statistical parametric maps of grey matter volume differences in subjects with High and Low Social Intelligence at p < 0.05, FWE cluster-level corrected.

## Discussion

The main goal of this study was to identify structural changes in the human brain associated with social intelligence, measured by the Guilford-Sullivan test. We observed that participants with high scores in the test demonstrated more GMV in the bilateral caudate, insula, IPL, ITG, and MOG. However, after applying conservative cluster-level correction, only findings in the bilateral caudate survived.

The cluster revealed in the present study occupies the head of the caudate nucleus. According to diffusion tensor imaging and retrograde tracing studies, this area is connected to the medial PFC (Lehéricy et al., 2004), which is considered to be responsible for emotional processing, decision-making, memory, self-perception, and social cognition in general (Bicks et al., 2015). Notably, the medial PFC is part of the TOM network (Carrington and Bailey, 2009; Dufour et al., 2013; Molenberghs et al., 2016). Structural connections with regions of the TOM system suggest the involvement of the caudate nucleus in brain mechanisms of social cognition.

This notion is supported by structural, functional, and clinical studies on brain mechanisms of social cognition. A genome-wide meta-analysis identified a positive correlation between RMET scores and volumes of the caudate nucleus and putamen (Warrier et al., 2018). The activity of the caudate nucleus, among other areas, was revealed during observing animations of mental interactions in the Triangles Task (Martin et al., 2016), during joint attention (induced by video stimuli) (Williams et al., 2005), and during guessing the mental state of a subject featuring first-person perspective sentences (Otsuka et al., 2011). In addition, several studies associated the activity in the caudate with understanding and classifying verbal expressions, which can be regarded as the verbal aspect of the TOM ability (corresponding with the social translations subtest in the Guilford-Sullivan test) (Shibata et al., 2010). A clinical study also demonstrated that focal left caudate nucleus lesions lead to impairment in TOM-related social cognition: poorer performance in the Faux-Pas test, the RMET, the Emotion Recognition Test (particularly for facial expressions of fear and sadness), accompany signs of alexithymia, social anhedonia, and reduced empathy abilities (J. Kemp et al., 2013). In another study, interpersonal schizotypy, characterised by impairments in social processes, was significantly negatively correlated with activation in the bilateral precuneus and right caudate nucleus during mentalising in the prisoner's dilemma game (Acosta et al., 2019). Thus, the results of structural and functional studies on healthy persons and patients as well as our own results indicate that the caudate nucleus participates in brain mechanisms of TOM-related socio-cognitive processes.

It is important to note the role of the caudate nucleus in brain mechanisms of socially oriented behaviour (e.g., trust and cooperation). Its activity was increased during signalling and integrating reputations gained through experience into trust decisions (Wardle et al., 2013). Being faced with choices, whether cooperative or not, was associated with activity in the medial PFC, anterior cingulate gyrus, caudate, and insula (Lemmers-Jansen et al., 2018). Specifically, cooperative choices were associated with activity in the right parietal cortex and caudate. A similar study on first-episode psychosis patients (characterized by reduced TOM abilities) showed reduced activation of the caudate and medial PFC during cooperative choices (Lemmers-Jansen et al., 2019). Another study on patients with early psychosis confirmed reduced caudate activation during cooperative actions (Fett et al., 2019). Therefore, the caudate nucleus demonstrates involvement in trust and cooperation through increased local neuronal activity and differential activity in patients and healthy subjects.

Moreover, changes in GMV, activation, and functional connectivity in the caudate were associated with impairments in social functioning. Some studies have found enlargement of the caudate in cases of autistic spectrum disorder (ASD), bipolar affective disorder, and early-onset schizophrenia, which are characterised by social behaviour impairments (Juuhl-Langseth et al., 2012; Ong et al., 2012; Qiu et al., 2016). In addition, a meta-analysis of atypical emotional face processing found ASD-related hyperactivation in the bilateral thalamus and bilateral caudate (Aoki et al., 2015). Furthermore, functional connectivity MRI studies have revealed less pronounced or absent connectivity between caudate nuclei and the cerebral cortex in ASD patients than in healthy controls (Turner et al., 2006). In conclusion, there is strong evidence of the association between social cognition alterations, which are due to pathologies and characteristics of the caudate nucleus.

In summary, the results of previous neuroanatomical and neurophysiological studies on healthy persons and patients provide evidence for the implication of the caudate nuclei in brain mechanisms of social cognition. However, this region was not revealed in meta-analytic studies assessing TOM-related experimental tasks. Thus, our findings of a relationship between the GMV in the caudate nuclei and the Guilford-Sullivan test of social intelligence scores can potentially be explained by the difference in structural characteristics of this area depending on the level of social intelligence.

While findings in the left insula did not survive conservative cluster-level thresholding, it was the second largest cluster. We suggest that it may play a role in neural mechanisms of social intelligence. For example, the anterior insular cortex is known to be the core element of empathy, which is closely related to affective TOM (Fan et al., 2011). In addition, activations in the insula were observed during pain judgement in TOM-measuring tasks (Corradi-Dell’Acqua et al., 2014). In addition, it was one of the regions implicated in decision making on whether to be cooperative or not (Lemmers-Jansen et al., 2018). Interestingly, the activation in both regions, revealed in the current study (the bilateral caudate and left insula), was associated with romantic love, which is an important social behaviour. Right caudate activation was correlated with the intensity of romantic passion, and left insula-putamen-globus pallidus activation was correlated with trait affect intensity (Aron et al., 2005). Importantly, there is evidence of involvement of the insula in pathologies associated with social cognition deficits. In schizophrenia patients, its activation was positively correlated with social loneliness (Lindner et al., 2014), while a meta-analysis of neuroimaging studies of social functions showed, compared to healthy controls, hypoactivation of the right anterior insula in ASD patients (Di Martino et al., 2009). In another study, reduced connectivity between the insula and brain regions involved in emotional and sensory processing was observed in patients with ASD (Ebisch et al., 2011). A VBM study has also revealed the decreased volume of insular subregions in cases of social anxiety disorder (Kawaguchi et al., 2016).

Thus, the implication of the insula in neural mechanisms of socio-cognitive processes is reflected by its structural and functional characteristics as well as by differential involvement in patients. Therefore, although findings in the insula did not survive conservative cluster-level thresholding in the current study, further research is needed to clarify its implication in social intelligence.

Our VBM analysis did not reveal regions reported in clinical studies that utilized Guilford-Sullivan test (cingulate cortex and orbitofrontal cortex). This can be explained by the fact that findings revealed by studying pathologies do not always extend to healthy people. Our experiment also did not find GMV changes in brain structures, attributed to the TOM neural system, in particular, the medial PFC, the TPJ in both hemispheres, the precuneus, and the right superior temporal sulcus. There are several reasons potentially explaining this fact. First, to our knowledge, there are no studies on the correlation between performances in TOM-related tests and the Guilford-Sullivan test. The latter is a specific measure of social intelligence. This ability and TOM, although closely related to each other, are not identical and, therefore, can be associated with different brain structures.

Second, a recent meta-analytic study demonstrated that although there are common regions shared across all TOM tasks, different types of TOM tasks reliably elicit activity in unique brain areas and are associated with distinct neural systems (Molenberghs et al., 2016). The authors found dissociation based on tasks with different instructional focuses (implicit versus explicit TOM tasks), types of mentalising inferences (cognitive versus affective TOM tasks), and modalities of presentation (visual versus verbal TOM tasks). For example, explicit tasks elicited more activation in the posterior medial frontal cortex and left TPJ, while implicit tasks elicited significantly greater activation in the dorsal medial PFC and right lateral orbitofrontal cortex. Accordingly, the Guilford-Sullivan test can be viewed as a specific type of TOM task or a more sophisticated task, measuring multiple components of TOM. Therefore, it can be associated with unique brain structures.

Studies that most accurately defined the nodes of the TOM system (using meta-analysis or large data samples) were based on functional data (Dufour et al., 2013; Molenberghs et al., 2016), while this study utilized VBM analysis to estimate the brain’s structural characteristics. A possible assumption is that brain areas demonstrating their involvement in the execution of action through changing their functional activity are not the same as those involved by changing their GMV. On a similar note, recent studies have demonstrated the existence of the so-called hidden nodes of neural systems that demonstrate their involvement by changing functional interactions with other brain areas without changing their local activity (Medvedev et al., 2019).

In summary, the reason our hypothesis was not supported may lie in the type of task (the Guilford-Sullivan test) or in the applied methodology (the VBM analysis).

## Limitations

Despite its advantages, this study has some possible limitations. One of them is associated with the nature of the Guilford-Sullivan test. Although its authors claim the independence of social intelligence and general cognitive abilities, some studies have revealed the correlation between them for some subtests of the Guilford-Sullivan test (Riggio et al., 1991; Shanley et al., 1971). This problem can potentially be resolved by using the general intelligence level as an additional covariate in the VBM analysis. In addition, it could be advantageous to consider other measures, reflecting levels of empathy, altruism, or the tendency for prosocial behaviour (e.g., trust and cooperation), as analysis covariates in future research.

## Conclusion

This VBM study provides new data suggesting the role of the bilateral caudate in social intelligence by demonstrating its higher GMV in people with high, compared to low, scores on the Guilford-Sullivan test. According to this result and previous data, there is strong evidence for the involvement of the caudate nucleus in the process of social cognition. Considering its structural and functional characteristics, the caudate nucleus can serve as a node in the neural system, underlying socio-cognitive processes and, in particular, TOM-related brain processes. However, future investigation is needed to confirm this assumption.

## Acknowledgments

We thank Maria Starchenko for her help with obtaining and preparation of psychological data.

